# Asymmetries in hue measured behaviorally and with visual evoked potentials

**DOI:** 10.1101/2025.10.20.683397

**Authors:** Jesse R. Macyczko, Osman B. Kavcar, Michael A. Crognale, Michael A. Webster

## Abstract

Hue percepts vary more rapidly along some directions in color space (e.g. near yellow) than others (e.g. near green) with corresponding differences in the size or stimulus range of different hue categories. The basis for these differences is not known. We examined whether the asymmetries are present in early cortical color coding by comparing the strength of hue differences using visual evoked potentials (VEPs) recorded from occipital cortex. Stimuli were spatial gratings with a fixed nominal contrast in the cone-opponent plane that varied sinusoidally in hue rather than saturation. The responses to different levels of hue separation were measured by the amplitude of the frequency-tagged signals and also in behavioral measurements employing a contrast matching task. For both, the same separation in hue angle resulted in stronger responses for angular differences centered on the yellow quadrant of the cone-opponent space. Responses were also larger for the yellow than blue quadrant, ruling out a general sensitivity loss to the blue-yellow axis as the basis for the differences. The responses differences paralleled the asymmetries in the rates of change in color appearance based on analyses of previous measures of hue scaling functions. The presence of these asymmetries in the VEP responses suggests that they arise relatively early in the cortical sensory representation of color rather than emerging late as a product of inference or color category learning.

## Introduction

Despite the smooth and continuous variation of wavelengths across the visible light spectrum, the experience of colors in the spectrum is non-uniform and instead appears as a limited number of bands or categories. Many studies have addressed the nature of these color categories and whether they reflect inherent biological constraints or learned characteristics of the world (Lindsry and Brown, 2021; Witzel and Gegenfurtner, 2018). However, a question that has received less attention is why these categories differ dramatically in size, or in the range of spectral stimuli they encompass. These differences are evident in many different representations of chromatic stimuli. For example, in the rainbow, red, green, and blue hues span a wide set of wavelengths while the yellow band is conspicuously narrow (Campbell, 1986). When hues are instead represented in terms of the cone receptor signals, rather than the physical stimulus, the size of different hue categories similarly vary widely. For example, common cone-opponent spaces define the stimuli in terms of the opposing signals in the long vs medium wavelength cones (LvsM) or signals in the short wavelength cones opposed by L and M (SvsLM). These dimensions correspond to the two cardinal axes of color coding in cells in the retina and geniculate (Derrington, Krauskopf and Lennie, 1984). Yet within this space, the region experienced as yellow hues again corresponds to only a narrow range of angles, while categories like green and purple extend across a wide sector (Figure 1). These differences can be quantified in hue scaling functions, which measure the proportion of red vs green or blue vs yellow perceived in the stimulus. The scaling functions vary only gradually along greenish or magenta angles in the cone-opponent plane (De Valois et al. 1997; Malkoc, Kay and Webster, 2005; Emery et al. 2017a) or across the wavelength spectrum (Gordon, Abramov, and Chan, 1994) but change more rapidly near yellowish hues. Moreover, the thresholds for discriminating changes in hue angle are also lower within the bluish and yellowish quadrants of the cone-opponent space, compared to the corresponding magenta and green quadrants (Krauskopf and Gegenfurtner, 1992; Hedjar, Toscani, and Gegenfurtner, 2025). Minima in discrimiation thresholds around bluish and yellowish-orange hues are also evident in wavelength discrimintation functions (Abramov, Gordon, and Chan, 2009; Krudy and Ladunga, 2001). Finally, the differences in category extent also persist even in spaces designed to equate perceptual differences in color. For instance, color naming in the Munsell system again reveals only a small range of hue values labeled as yellow, compared to wider swaths for blue and green and red (Lindsey and Brown, 2006).

**Figure 1.**
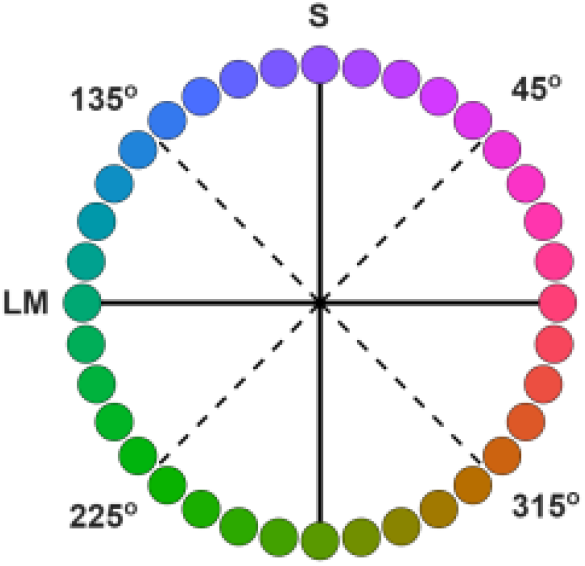
Color variations within the LvsM and SvsLM cone-opponent plane illustrating the differences in the range of hue angles for different color categories. Depicted colors are only approximate and for illustration purposes only.

The basis for the size differences between color categories is not understood (Witzel and Gegenfurtner, 2018). The fact that they are seen in stimuli are matched for the independent LvsM and SvsLM signals (as in Figure 1) suggests that they cannot be explained by how the color signals represented within the cardinal mechanisms of the retina and geniculate (De Valois et al., 1997). However, how and where they might arise at cortical levels remain unknown. In this study, we asked if they are present in early cortical stages of color processing by examining the rate of hue changes at different regions in the cone-opponent plane using visual evoked potentials (VEPs) (Kulikowski et al. 1987; Rabin et al. 1994). The VEP measured at the occipital pole is thought to be primarily driven by neural responses in primary visual cortex and early extrastriate areas, and for example, is unaffected by cortical color losses (dyschromatopisa) arising from damage at later stages (Victor et al., 1989; Crognale et al. 2013).

Chromatic VEPs are also surprisingly impervious to attentional effects (Highsmith & Crognale, 2010; Wang & Wade, 2011; Arthur et al., 2024) and thus are less likely to be susceptible to cognitive or linguistic effects that have been found to influence categorical processing of color (Witzel and Gegenfurnter, 2018). Thus, the VEPs provide a potential test for whether the hue asymmetries reflect the low-level sensory representation of color rather than potentially higher-level interpretations or learned associations in the formation of color categories.

To explore these questions, we measured the strength of hue changes within the four color quadrants of cone-opponent space of Figure 1, for hue angles centered at 45 (magenta), 135 (blue), 225 (green), and 315 (yellow) degrees (as illustrated in Figure 1). The stimuli were equiluminant sinusoidal gratings, but instead of varying in contrast or saturation, were modulated in hue angle at a constant chromatic contrast. The “contrast” in the stimuli therefore corresponded to the magnitude of the hue angle differences and increased as the colors separated away from each other. This allowed us to generate “hue” contrast response functions (CRFs) for the response amplitudes as a function of the hue angle difference. Finally, the same stimuli were also used in behavioral experiments which measured when the gratings in different quadrants appeared to match in contrast, as well as in reanalyses of color appearane changes based on prior studies of hue scaling functions (Emery et al., 2017a, 2023). This allowed us to examine whether regions of color space with rapid apparent changes in hues, such as the 315^°^ quadrant, would lead to steeper CRFs than regions with gradual changes in hues, such as the 225^°^ quadrant, and whether these effects were manifest in both the VEPs and perceptual judgment.

## Methods

### Participants

Ten students from the University of Nevada, Reno participated in the experiments (5 males and 5 females, average age of 28). All participants were naive to the aims of the experiments and had normal or corrected acuity and normal color vision as assessed by Cambridge Colour Test (Cambridge Research Systems). Protocols were approved by UNR’s Institutional Review Board (IRB), and participation was with informed consent. All participants completed both the behavioral and VEP experiments. Results reported are based on the average results across the observers.

### Apparatus

Stimuli were generated using Psychtoolbox (Brainard, 1997) in Matlab (Mathworks, USA) and displayed on a VIEWPixx / 3D monitor (VPixx Technologies Inc.) with a refresh rate of 120Hz. The monitor was calibrated using a PR-655 SpectraScan spectroradiometer (Photo Research). VEPs were recorded using gold-plated Grass electrodes with a single active site located at Oz, the reference point at Pz, and the ground electrode placed on the forehead (international 10-20 system) (Odom, 2016). Electrodes were fixed using conductive paste with impedance kept below 10 kΩ at 30Hz. Gross potentials were sampled using Grass P5 Series at 1,000Hz, amplified, then notch-filtered at 60Hz (Grass Instrument Co.). High pass and low pass analog filtering was set to 0.3Hz and 100Hz, respectively. The signals were relayed to a PC using a National Instruments input/output board and were digitally low pass filtered offline at 100Hz.

### Stimuli

Stimuli for both the behavioral and VEP experiments were based on a scaled version of the Derrington-Krauskopf-Lennie (1984) space, which represents stimuli in terms of the LvsM and SvsLM signals relative to a neutral gray level. For our space this gray had a chromaticity equal to Illuminant D65 and cone spectral sensitivities based on Stockman and Sharpe (Stockman, 2019). Units along the two axes were scaled to roughly equate sensitivity to contrasts along the two axes. The resulting stimulus coordinates are related to the l_mb_,s_mb_ coordinates of the MacLeod-Boynton (1979) chromaticity diagram by:

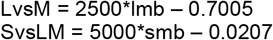

Where 0.7005, 0.0207 are the coordinates of the neutral gray. Hue angles and contrasts correspond to the polar coordinates of the stimuli. Stimuli were always centered along one of the 4 diagonals of the plane, at 45^°^, 135^°^, 225^°^, and 315^°^ (Figure 1). For each of these quadrants spatial gratings were generated that varied sinusoidally between two hue angles offset equally from the axes (Figure 2). The separation between the hue angles varied from 0^°^ to 100^°^ of in steps of 20^°^. For example, a grating in the 135^°^ quadrant with a hue separation of 100^°^ varied in hue angles from 85^°^ to 185^°^ in the plane. The chromatic variations were shown at a constant luminance of 20 cd/m^2^ (with equiluminance determined for individual observers by a minimum motion task (Cavanagh, MacLeod and Anstis, 1987) and a constant chromatic contrast of 80 units in the cone-opponent space as specified above. The gratings had a spatial frequency of 1 c/deg and presented as full-field stimuli, subtended 53^°^ by 30^°^ at the viewing distance 57cm.

**Figure 2.**
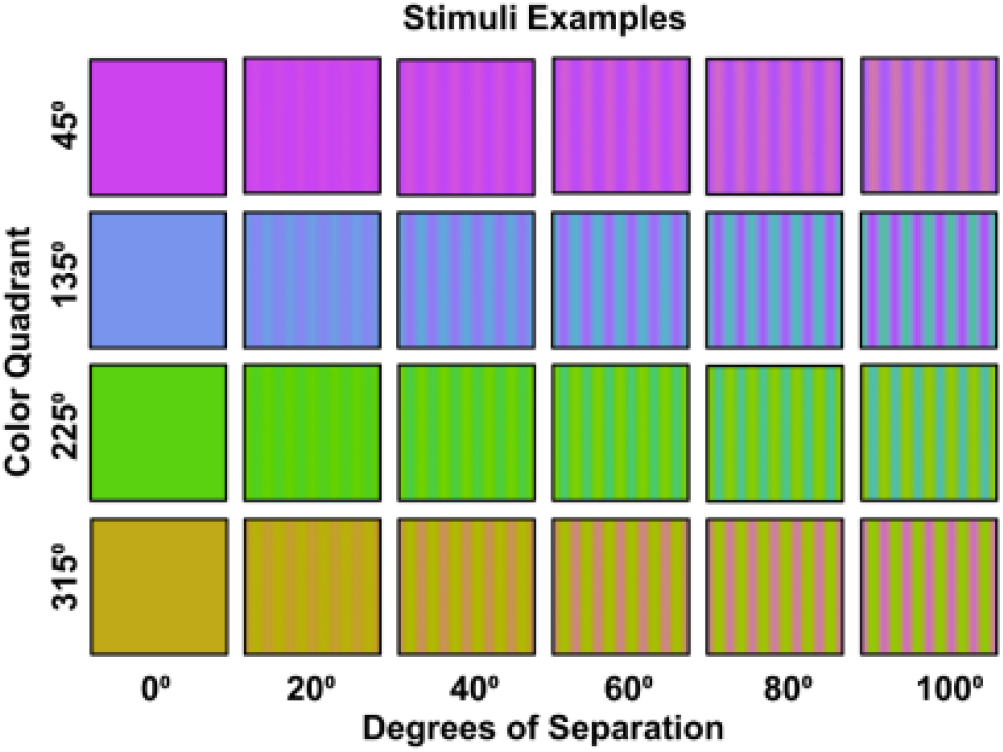
Examples of the hue gratings for different angular separations at each of the four color quadrants.

### Behavioral Measures - suprathreshold color difference matching

We compared the perceptual magnitude of hue differences across the four color quadrants using a suprathreshold matching task (Switkes & Crognale, 1999). In this task, participants were asked to adjust the difference between the two hues in a test grating from one quadrant to match the difference between two hues in a reference quadrant. The two stimuli were presented at the same time, vertically stacked with the reference quadrant on top, for 50ms, chosen to match the stimulus duration for the VEP stimuli. The presentation was followed by a gray screen until subjects provided their response. This adjustment was done by increasing or decreasing the degrees of separation between hues in the test quadrant using a staircase method with 13 reversals (with the matching value based on the mean of the final 10 reversals). The 225^°^ quadrant was fixed as the reference grating while the 45^°^, 135^°^, and 315^°^ quadrants were used as the test gratings. Matches were made for reference gratings set to 20^°^, 40^°^, 60^°^, 80^°^, and 100^°^ of hue angle separation. These were shown in random order for each test quadrant. Participants completed matches for each of the three test quadrants three times, in counterbalanced order (run one order: 315^°^, 135^°^, 45^°^; run two order: 45^°^, 135^°^, 315^°^; run three order: 135^°^, 315^°^, 45^°^), with the mean matches based on the mean of the three repeated settings for each test angle and contrast.

### VEP

Neural responses were measured with a steady state VEP paradigm, which tracks the responses at the stimulus temporal frequency and has the advantage of rapid estimates and high signal to noise (Norcia et al. 2015). We compared VEP amplitudes at the presentation rate in response to the hue differences across the four color quadrants. The VEPs were measured for the same hue different steps as in the behavioral measures but in addition included a zero-contrast difference, so that there were 6 separations of 0^°^, 20^°^, 40^°^, 60^°^, 80^°^, and 100^°^. For each quadrant, the six levels of hue separation were presented in random order, with each level being shown for 50 seconds. Participants were able to take breaks between levels. Participants completed each of the four quadrants two times, again counterbalancing for order (run one: 135^°^, 45^°^, 225^°^, 315^°^; run two: 225^°^, 315^°^, 135^°^, 45^°^). Gratings were presented for 50ms and repeated at 2Hz (Figure 3). The gratings were followed by a uniform field with the average color (i.e. zero separation) for 200ms and then alternated with a uniform field of the average complementary color for 250ms. This alternation was included to control for chromatic adaptation during the session, which could shift the mean cone responses toward gray and thus alter the hue relations in the stimuli. The complementary chromaticity instead ensured that the chromatic adaptation was equal to the gray background for all stimuli for the presentation rates used (Webster and Wilson, 2000).

**Figure 3.**
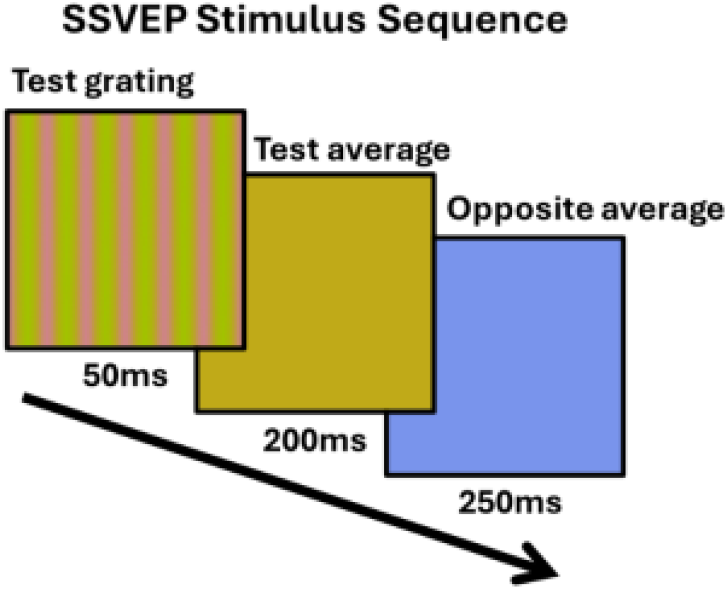
Illustration of the stimulus sequence during the SSVEP recordings. The test grating is first presented for 50ms, followed by the average color of the test grating for 200ms, then lastly the opposite hue angle of the average color. This was done to avoid chromatic adaptation.

Responses were averaged across trials for each participant. The amplitudes of the responses at the first four unique harmonics of the stimulus frequency (2Hz, 6Hz, 10Hz, and 14Hz) were computed using fast Fourier transform and summed after subtracting the surrounding noise. This noise was computed by averaging the surrounding 20 bins of the target bin, corresponding to 0.4Hz (Retter et al., 2021).

## Results

### Behavioral Measures

Results for the matching experiment are shown in Figure 4, which plots the angular differences within each quadrant that appeared to match a fixed angular difference in the “green” (225^°^ quadrant). The matches suggest that smaller differences were required within each of the three comparison quadrants. This was confirmed statistically by one-sample t-tests comparing the differences between the test and reference quadrants against the null of no difference across the different levels of angular separation.

**Figure 4.**
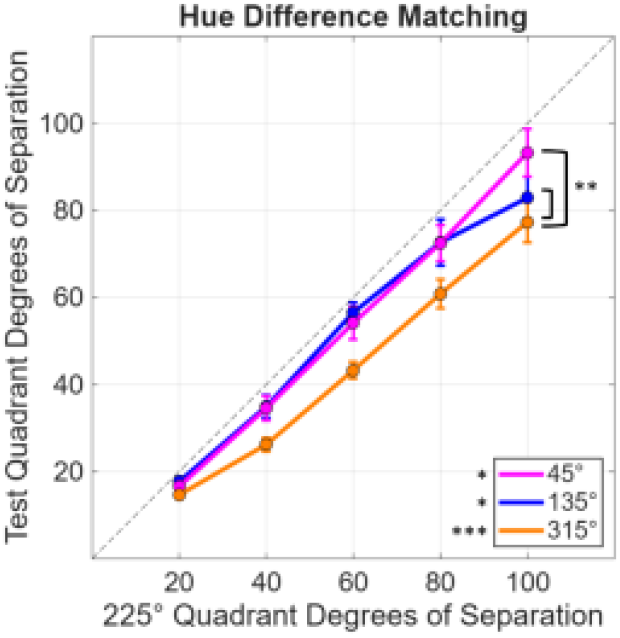
Hue separation within each test quadrant required to match a fixed separation in the 225^°^ reference quadrant. Significant statistical tests are labeled with *(*p* < 0.05) or ***(*p* <.001). Asterisks by the line plots indicate repeated-measures two-way ANOVA pairwise tests while those beside the legend indicate one-sample t-tests. Error bars indicate 1 SEM.

Specifically, the differences between the test and reference quadrants for each contrast separation were averaged together, giving an overall difference for each of the test quadrants. The mean difference for the 45^°^ quadrant was -5.79 (sd = 9.36; t(9) = -1.96, p =.041); for the 135^°^ quadrant 7.06 (sd = 8.98; t(9) = -2.49, p =.017); and for the 315^°^ quadrant 15.55 (sd = 6.96; t(9) = -7.07, p <.001; after applying the Benjamini–Hochberg (1995) correction (Q values of 0.05) for multiple comparisons).

To compare the differences among the 3 comparison quadrants, a repeated-measures two-way ANOVA was used (3 quadrants by 5 contrast levels). There were significant main effects of quadrant (*F*(2, 18) = 13.47 *p* <.001) and hue separation (*F*(4, 36) = 232.98, *p* <.001) with a significant interaction (*F*(8,72) = 3.54, *p* =.002). The interaction is likely driven by the convergence of the curves at the lower contrasts. Follow up pairwise comparisons using Benjamini-Hochberg correction indicated that the 315^°^ quadrant was significantly lower (or required a smaller separation in contrast to match the reference 225^°^ quadrant) than both the 45^°^ (p =.003) and the 135^°^ (p =.002) quadrants.

### VEP

Responses at the 2Hz presentation rate increased systematically with the stimulus angle difference and are well fit by a Naka-Rushton (1967) contrast response function (CRF) (Figure 5). Comparisons across the quadrants were restricted to the amplitudes at 40-100^°^ degree of separation to ensure that responses were above threshold. Visual inspection of the curves suggests that increasing separation of hue angles in the 315^°^ quadrant led to a more rapid increase in SSVEP amplitude compared to the 45^°^, 135^°^, and 225^°^ quadrants. This was confirmed in a repeated-measures two-way ANOVA, which showed significant main effects of quadrant (*F*(3, 27) = 7.11 *p* =.001) and hue separation (*F*(3, 27) = 46.26, *p* <.001) with a significant interaction (*F*(9,81) = 2.99, *p* =.004). As was the case in the behavioral experiment, the interaction is likely driven by the convergence of the curves at the lower contrasts where the signals were approaching threshold. Follow up comparisons using Benjamini-Hochberg correction (Q values of 0.05) indicated that the 315^°^ quadrant was higher than the 45^°^ (p <.001), 135^°^ (p =.03), and 225^°^ (p =.03) quadrants. Note that the stronger responses in the315^°^ quadrant are despite the weaker responses to the average hues shown as uniform fields (0^°^ separation), which are instead consistent with the weaker sensitivity to the blue-yellow axis in cortical responses (Goddard et al., 2010).

**Figure 5.**
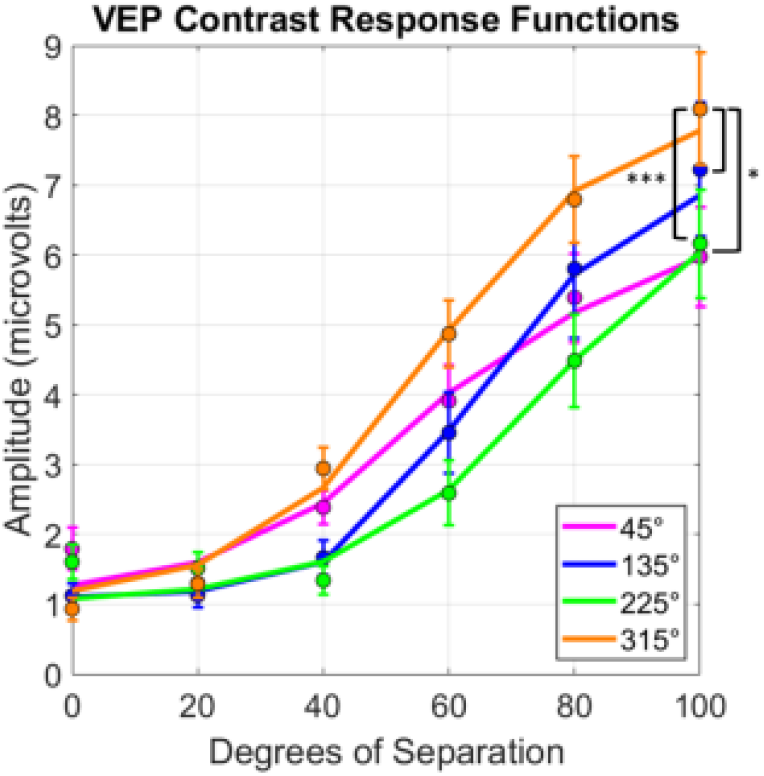
VEP responses (at the 2 Hz presentation frequency and summed harmonics) to increasing separation of hues across the four color quadrants. Dots indicate averages and lines indicate fitted CRFs. Significant pairwise comparisons are labeled with *(*p* < 0.05) or ***(*p* <.001) after Benjamini-Hochberg corrections. Error bars indicate 1 SEM.

### Relation to hue scaling functions

As noted in the Introduction, our study was motivated by the common observation that hue percepts vary at different rates in different regions of the wavelength spectrum or different angles in color space. One potential account of the observed hue contrast responses in Figures 4 and 5 is thus that the stimuli match when the hue differences rather than the stimulus differences are equated. To explore this, we used the hue scaling functions of Emery et al. (2023) to predict when the stimulus differences along the 4 quadrants should match, based on the observers’ scaling functions. The average scaling function is shown in Figure 6a, which plots the “perceptual angle (based on the proportion of red vs. green or blue vs. yellow reported for the color) as a function of the stimulus angle (or direction within the cone opponent plane). Again, the function is not linear and instead varies more rapidly along some stimulus angles than others. This is illustrated in Figure 6b, which plots the derivative of the hue scaling function based on a polynomial fit. As before, we fixed the stimulus differences in the 225° (“green”) cone-opponent quadrant and then calculated the stimulus differences required to match the changes in perceptual angles along the remaining quadrants. The predicted matches are shown in Figure 6c, and replicate the general pattern of differences in both the contrast matching and VEP responses. In particular, the hue variations are again more rapid within the yellow quadrant compared to all three of the other quadrants, while the blue quadrant in this case is very similar to the green. We did not conduct quantitative comparisons of the hue scaling predictions with the quadrant differences measured in the present study because the stimuli and observers in the study were different. However, this analysis suggests that the differences measured in the cortical responses are qualitatively consistent with the color appearance differences measured by hue scaling.

**Figure 6.**
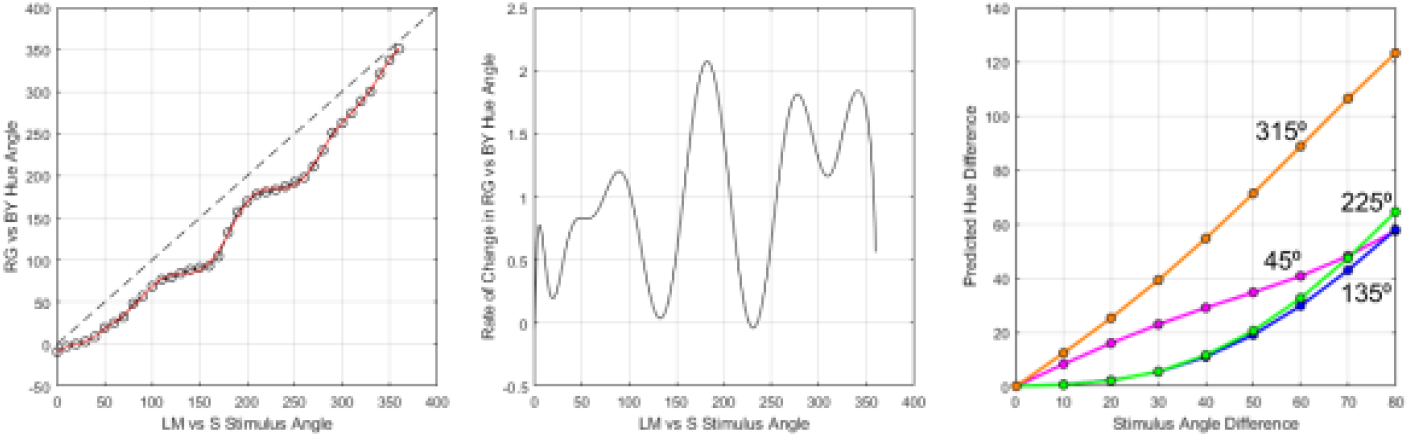
a) average hue scaling function from Emery et al. (2023), representing the hue of the stimulus as angle within a red vs green / blue vs. yellow plane, as a function of the stimulus angle in the LvsM and SvsLM cone-opponent plane. b) Rate of change in hue as a function of stimulus angle, based on the derivative of a polynomial fit of the hue scaling function. c) magnitude of hue differences (in perceptual angle) as a function of the difference in stimulus hue angle long each of the quadrants.

## Discussion

To summarize, we found that nonuniformities in the rate of hue changes (Figure 6) are mirrored in both the perceptual salience of hue differences (Figure 4) and the magnitude of neural responses measured with VEPs (Figure 5). To the extent that the occipital VEP reflects activity originating in early visual cortex, these parallels suggest that the asymmetries in hue percepts arise relatively early in the cortical processing of color.

VEPs have been used extensively to characterize color coding in the cortex, at both early and later levels, for example to explore the role of color in object processing (Martinovic et al. 2008; Retter et al. 2021). In early cortex, the responses point to a number of important transformations of the retino-geniculate inputs. First, The VEP waveforms for chromatic and luminance stimuli have distinct signatures, suggesting that they reflect different neural populations (Rabin et al. 1994). Responses to chromatic stimuli also show bandpass tuning, indicating that they are not only color-selective but also spatially selective. This spatial tuning is consistent with the emergence of double-opponent receptive fields in V1 (Shapley, Nunez, and Gordon, 2019). Other subpopulations have lowpass tuning and may reflect single-opponent or input layers to the cortex (Rozman et al. 2025). The VEPs we measured are presumably driven primarily by the double-opponent cells, because the responses to the grating stimuli were stronger than the responses to the uniform field (corresponding to the 0 difference in hue) (Figure 5). Moreover, the mean chromatiity of the test grating and it’s complementary color shifted toward gray as the hue angles composing the grating increased. This again suggests that the increasing responses with hue angle separation were driven by the increasing spatial contrast in the grating rather than by the contrast modulation in the mean color of the stimuli.

There are also large transformations in how the dimension of color itself is represented. In the retina and geniculate chromatic signals are largely mediated by the two cardinal axes carried along the parvocellular (LvsM) and koniocellular (SvsLM) pathways. However, multiple lines of evidence indicate that these signals are combined in V1 to form higher-order mechanisms tuned to intermediate directions in the cone-opponent plane. Multiple color mechanisms are also evident in the VEP responses to different directions in color space (Duncan et al. 2012; Chen and Gegenfurtner, 2021; Rozman et al. 2025; Shapley, Nunez, and Gordon, 2025). Our results also point to this representation, because, as noted, the differences we found between the four quadrants cannot be accounted for by separable responses within the independent LvsM and SvsLM axes.

Higher-order mechanisms for color are similar to how the cortex represents orientation, where the cells have a wide range of preferred angles rather than simply two canonical axes like horizontal and vertical. In population codes of this kind, features like orientation or hue (the “angle” in color space) may be represented by which cells are responding (e.g. by the peak of the response distribution) rather than by the relative responses within a small number of mechanisms (e.g. the cardinal or perceptual opponent axes, as in the Hering model of color opponency). The cortical code for color may therefore reflect a general design principle, and may be further enhanced by the increasing hue-selectivity of cells at later cortical stages (Zaidi and Conway, 2019). On the other hand, how the population responses are interpreted could be different. For example, Emery et al.(2023) showed that individual differences in the hue scaling functions reflected multiple independent factors narrowly tuned to different hues, whereas analogous measurements of “motion scaling functions” pointed to a representation referenced to the vertical and horizontal axes of space. This could reflect differences in how information is decoded from the responses, even if the distribution of responses themselves (in terms of the number and selectivity of the encoding mechanisms) is similar. For example, the readout for motion could (and may need to) retain the intrinsic coordinates of space, while different colors may be represented more like qualitatively different objects.

This distinction raises the question of whether the differences in hue across the different quadrants are due to how the color information is encoded vs. decoded. In other words, is the range of hues perceived as yellow compared to green a reflection of asymmetries in the population of color mechanisms, or in how those population responses are analyzed? As noted in the Introduction, asymmetries in color coding can arise very early. For example, wavelength discrimination functions vary across the spectrum in ways that can be precisely modeled by the cone absorption sepctra and how they mightcould be compared in early post-receptoral combinations (Kaiser and Boynton, 1996). Conversely, high-level influences on color perception and categorization are well established (Jameson and Andrade, 1997; Witzel and Gegenfurtner, 2018). Moreover, color categories, which as we noted seem to roughly track the ranges seen for different hue percepts, have been found to be lacking in macaque monkeys (Garside et al., 2025, despite sharing very similar color encoding stages to humans (De Valois et al., 1974; Neitz and Neitz, 2017). This suggests that the categories may be more a reflection of color concepts and language rather than sensory partitions, and thus consistent with decoding. However, the VEP responses presumably reflect the magnitude of the population responses, not how they are interpreted. This suggests that the hue differences may already arise from asymmetries built into the early cortical sites of encoding. Thus even if how they are categorized arises later, the population code may already have biases in the representation. (Another possibility is that the biases reflect feedback from higher areas, but are still manifest as differences on the population responses.) The VEP responses we recorded provide only a univariate overall measure of the cortical activity, and thus the response changes could be modulated in two different ways: by increasing the responses within the same cells, or by increasing the activity in the number of cells. We consider each of these in turn.

One well-established bias in color coding is that sensitivity is weaker along the blue-yellow axis of the cone-opponent plane (Webster, 2020), or alternatively along a “warm-cool” (orange to teal) direction (Manalansan, Whitehead, and Webster, 2025). This has been attributed to the higher variance of both illuminant and surface colors along these axes, so that vision might be more adapted and thus less sensitive to these axes. This predicts that neural responses should be stronger along the magenta and green (our 45° and 225° quadrants) than the blue and yellow directions (135° and 315°). In our experiments we nominally varied the hue difference between the stimuli, rather than the saturation. However, note that these hue differences correspond to contrast or saturation changes along the axes orthogonal to the quadrant. That is, when hues were varied relative to the yellow reference, the stimuli primarily alter contrasts along the color channels tuned to the magenta-greenish directions. Moreover, the signals along orthogonal directions show little masking or cross-talk, and thus should be relatively unaffected by the axis of the reference color (Chen and Gegenfurtner, 2021; Rozman et al, 2025). Thus the more robust responses around yellow could in part be because the responses in these quadrants are driven by mechanisms tuned to the magenta and greenish quadrants. Conversely, hue variations relative to the red and green reference quadrants might instead be detected by mechanisms tuned to the blue-yellow dimension, which again may have weaker sensitivity. One problem with this interpretation is that finer discrimination for hue angles around yellow cannot be accounted for by the weaker saturation along the yellow axis (Hedjar and Gegenfurtner, 2025). Moreover, the general loss in sensitivity for the bluish-yellowish axis also does not account for the large differences we found between the yellow and blue quadrants. However, there are also a number of asymmetries between these complementary color directions. Studies of color constancy have shown that observers are less sensitive to changes in bluish than yellowish illuminants (Hurlbert and Yu, 2025). Moreover, bluish tints are more likely to be attributed to the lighting while equivalent yellowish chromatic contrasts are more likely to be perceived as a surface reflectance (Winkler et al. 2015). Finally, background colors tend to be more greenish and blue, or cool colors, while objects have been found to be more strongly associated with warm colors (Rosenthal et al., 2018). Neural responses to color also tend to be stronger for warm colors, though this bias emerges more clearly at later cortical sites (Rosenthal et al. 2018, 2021; Pennock et al. 2023), and even beginning in the retina there are asymmetries in the properties of neurons that excited (S-ON) or inhibited (S-OFF) by signals from the S cones (Martin, 2023). In this regard, another important transformation in cortical color coding is the emergence of unipolar chromatic responses, which could arise from half-wave rectification of the responses and is consistent with the low spontaneous activity of cortical cells (so that they can signal excitation but not directly inhibition) (von der Twer and MacLeod, 2001). These unipolar mechanisms may also be more characteristic of the spatiochromatic mechanisms reflected in the the VEP responses to spatial color patterns (Rozman et al., 2025), Thus collectively these findings point to potential biases in the population responses to yellow vs. blue.

At the level probed by occipital VEPs, these biases do not appear to reflect differences in the tuning or selectivity for color, because this has been found to be broad and largely independent of color direction (Chen and Gegenfurtner, 2021; Rozman et al. 2025; though the chromatic tuning narrows substantially at higher cortical stages; Zaidi and Conway, 2019). Tuning functions revealed by chromatic contrast adaptation also are consistent with broad linear mechanisms biased along the cardinal axes (Krauskopf, Williams, and Helley, 1982; Webster and Mollon, 2024), though this adaptation may include a strong component from the response changes in early cortical stages such as the cardinal mechanism inputs in V1 (Tailby et al., 2005). Instead, the stronger hue responses around yellow could instead reflect the higher density of mechanisms tuned around this direction. By this account, the activity for hue contrasts relative to the yellow would include more cells tuned to color directions within the yellow quadrant compared to the remaining quadrants. In fact a number of studies have reported a preponderance of cells tuned to blue-yellow directions in the cortex (Conway, 2001; Horwitz, et al., 2007; Lafar-Sousa and Conway, 2013; Solomon and Lennie, 2005). This higher density could in turn be driven by the higher significance of the daylight axis (Horwitz, 2020) or of warm hues (Rosenthal et al. 2018; Pennock et al. 2023) and the potential need to more precisely represent and discriminate between them. By this account, the differences in VEP responses could partly be driven by recruitment of a broader population of cells as the colors in the gratings increasingly differ, with the magnitude of these effects dependent on the relative biases in the cell population along different color axes.

A final set of questions is why these differences in response magnitudes lead to the changes in hue percepts, and why these biases are not “corrected” in the experience of color. One possibility is that the distortions in the population lead to distortions in the readout. For example, hue might vary according the variations in the population rather than the stimulus because of uniform sampling of a nonuniform distribution. That is, changes in hue might vary not according to equal steps in the stimulus space, but according to constant shifts in the population responses to the stimulus. Variations in hue may then be given more weight in regions of the color space where these differences are more important and thus sampled more finely. This would corrupt a faithful representation of hue in terms of the stimulus and corresponding cone excitations. However, unlike spatial dimensions such as orientation or motion, there may be little intrinsic value in a metrical representation for color (Emery et al., 2023), and thus less need to map the percepts into a stimulus-referenced coordinate frame. The differences between hues may instead reflect how important it is to judge those stimulus differences.

## Acknowledgments

Supported by EY-010834 and P30 GM145646

## Notes

### Competing Interest Statement

The authors have declared no competing interest.

## References

Abramov, I., Gordon, J., & Chan, H. (2009). Color appearance: Properties of the uniform appearance diagram derived from hue and saturation scaling. Attention, Perception, & Psychophysics, 71(3), 632–643.

Arthur C., Kavcar O.B., Wise M.V. & Crognale M.A. (2024). Pattern reversal chromatic VEPs like onsets, are unaffected by attentional demand. Visual Neuroscience, 41, E006. 10.1017/S0952523824000063

Benjamini, Y., & Hochberg, Y. (1995). Controlling the false discovery rate: a practical and powerful approach to multiple testing. Journal of the Royal statistical society: series B (Methodological), 57(1), 289–300.

Brainard, David H., and Spatial Vision. “The psychophysics toolbox.” Spatial vision 10.4 (1997): 433–436.

Campbell, F. W. (1986). In search of the spectrum’s elusive yellow. Ophthalmic and Physiological Optics, 6(2), 129–133.

Cavanagh, P., MacLeod, D. I., & Anstis, S. M. (1987). Equiluminance: spatial and temporal factors and the contribution of blue-sensitive cones. Journal of the Optical Society of America A, 4(8), 1428–1438.

Chen, J., & Gegenfurtner, K. R. (2021). Electrophysiological evidence for higher-level chromatic mechanisms in humans. Journal of Vision, 21(8), 12–12.

Conway BR. 2001. Spatial structure of cone inputs to color cells in alert macaque primary visual cortex (V-1). J. Neurosci. 21:2768–83

Crognale, M. A., Duncan, C. S., Shoenhard, H., Peterson, D. J., & Berryhill, M. E. (2013). The locus of color sensation: cortical color loss and the chromatic visual evoked potential. Journal of vision, 13(10), 15–15.

De Valois, R. L., De Valois, K. K., Switkes, E., & Mahon, L. (1997). Hue scaling of isoluminant and cone-specific lights. Vision research, 37(7), 885–897.

De Valois, R. L., Morgan, H. C., Polson, M. C., Mead, W. R., & Hull, E. M. (1974). Psychophysical studies of monkey vision—I. Macaque luminosity and color vision tests. Vision research, 14(1), 53–67.

Duncan, C. S., Roth, E. J., Mizokami, Y., McDermott, K. C., & Crognale, M. A. (2012). Contrast adaptation reveals the contributions from chromatic channels tuned to intermediate directions of color space in the chromatic visual evoked potential. Journal of Vision, 12(9), 63–63.

Emery, K. J., Volbrecht, V. J., Peterzell, D. H., & Webster, M. A. (2017a). Variations in normal color vision. VI. Factors underlying individual differences in hue scaling and their implications for models of color appearance. Vision research, 141, 51–65.

Emery, K. J., Volbrecht, V. J., Peterzell, D. H., & Webster, M. A. (2017b). Variations in normal color vision. VII. Relationships between color naming and hue scaling. Vision research, 141, 66–75.

Emery, K. J., Volbrecht, V. J., Peterzell, D. H., & Webster, M. A. (2023). Fundamentally different representations of color and motion revealed by individual differences in perceptual scaling. Proceedings of the National Academy of Sciences, 120(4), e2202262120.

Garside, D. J., Chang, A. L., Selwyn, H. M., & Conway, B. R. (2025). The origin of color categories. Proceedings of the National Academy of Sciences, 122(1), e2400273121.

Goddard, E., Mannion, D. J., McDonald, J. S., Solomon, S. G., & Clifford, C. W. (2010). Combination of subcortical color channels in human visual cortex. Journal of vision, 10(5), 25–25.

Gordon, J., Abramov, I., & Chan, H. (1994). Describing color appearance: Hue and saturation scaling. Perception & psychophysics, 56(1), 27–41.

Hedjar, L., Toscani, M., & Gegenfurtner, K. R. (2025). Importance of hue: color discrimination of three-dimensional objects and two-dimensional discs. Journal of the Optical Society of America A, 42(5), B296–B304.

Highsmith, J & Crognale M.A. (2010). Attentional shifts have little effect on the waveform of the chromatic onset VEP. Ophthalmic and Physiological Optics, 30, 525–533

Horwitz, G. D. (2020). Signals related to color in the early visual cortex. Annual review of vision science, 6(1), 287–311.

Horwitz GD, Chichilnisky EJ, Albright TD 2007. Cone inputs to simple and complex cells in V1 of awake macaque. J. Neurophysiol. 97:3070–81

Hurlbert, A., & Yu, C. (2025). Seeing the Light: Perception and Discrimination of Illumination Color. Annual Review of Vision Science, 11.

Jameson K.A., D’Andrade R.G. (1997) in Color Categories in Thought and Language, eds Hardin CL, Maffi L (Cambridge Univ Press, Cambridge, U.K.), pp 295–319

Kaiser, P. K., & Boynton, R. M. (1996). Human color vision. Optical Society of America: Washington DC.

Krauskopf, J., & Gegenfurtner, K. (1992). Color discrimination and adaptation. Vision research, 32(11), 2165–2175.

Krauskopf, J., Williams, D. R., & Heeley, D. W. (1982). Cardinal directions of color space. Vision research, 22(9), 1123–1131.

Krudy, A., & Ladunga, K. (2001). Measuring wavelength discrimination threshold along the entire visible spectrum. Periodica Polytechnica Mechanical Engineering, 45(1), 41–48.

Kulikowski, J. J., Murray, I. J., & Parry, N. R. (1987). Human visual evoked potentials to chromatic and achromatic gratings. Clin. Vis. Sci, 1, 231–244.

Lafer-Sousa R, Conway BR. 2013. Parallel, multi-stage processing of colors, faces and shapes in macaque inferior temporal cortex. Nat. Neurosci. 16:1870–78

Lindsey, D. T., & Brown, A. M. (2006). Universality of color names. Proceedings of the National Academy of Sciences, 103(44), 16608–16613.

Lindsey, D. T., & Brown, A. M. (2021). Lexical color categories. Annual Review of Vision Science, 7(1), 605–631.

MacLeod, D. I., & Boynton, R. M. (1979). Chromaticity diagram showing cone excitation by stimuli of equal luminance. Journal of the Optical Society of America, 69(8), 1183–1186.

Malkoc, G., Kay, P., & Webster, M. A. (2005). Variations in normal color vision. IV. Binary hues and hue scaling. Journal of the Optical Society of America A, 22(10), 2154–2168.

Manalansan, J., Whitehead, L. A., & Webster, M. A. (2025). Warm versus cool colors and their relation to color perception. Journal of Vision, 25(4), 13–13.

Martin, P. R. (2023). The Verriest Lecture: Pathways to color in the eye and brain. Journal of the Optical Society of America A, 40(3), V1–V10.

Martinovic, J., Gruber, T., & Müller, M. M. (2008). Coding of visual object features and feature conjunctions in the human brain. PloS one, 3(11), e3781.

Naka, K. I., & Rushton, W. A. H. (1967). The generation and spread of S-potentials in fish (Cyprinidae). The Journal of Physiology, 192(2), 437.

Neitz, J., & Neitz, M. (2017). Evolution of the circuitry for conscious color vision in primates. Eye, 31(2), 286–300.

Norcia, A. M., Appelbaum, L. G., Ales, J. M., Cottereau, B. R., & Rossion, B. (2015). The steady-state visual evoked potential in vision research: A review. Journal of vision, 15(6), 4–4.

Odom, J. V., Bach, M., Brigell, M., Holder, G. E., McCulloch, D. L., Mizota, A., … & International Society for Clinical Electrophysiology of Vision. (2016). ISCEV standard for clinical visual evoked potentials:(2016 update). Documenta Ophthalmologica, 133(1), 1–9.

Pennock, I. M. L., Racey, C., Allen, E. J., Wu, Y., Naselaris, T., Kay, K. N., Franklin, A., & Bosten, J. M. (2023). Color-biased regions in the ventral visual pathway are food selective. Current Biology, 33(1), 134–146.e4. 10.1016/j.cub.2022.11.063

Rabin, J., Switkes, E., Crognale, M., Schneck, M. E., & Adams, A. J. (1994). Visual evoked potentials in three-dimensional color space: correlates of spatio-chromatic processing. Vision research, 34(20), 2657–2671.

Retter, T., Rossion, B., Schiltz, C. (2021). Harmonic Amplitude Summation for Frequency-tagging Analysis. J Cogn Neurosci, 33 (11): 2372–2393. doi: 10.1162/jocn_a_01763

Retter, T. L., Gao, Y., Jiang, F., Rossion, B., & Webster, M. A. (2021). Early, color-specific neural responses to object color knowledge. BioRxiv, 2021–02.

Rosenthal, I., Ratnasingam, S., Haile, T., Eastman, S., Fuller-Deets, J., & Conway, B. R. (2018). Color statistics of objects, and color tuning of object cortex in macaque monkey. Journal of vision, 18(11), 1–1.

Rosenthal, I. A., Singh, S. R., Hermann, K. L., Pantazis, D., & Conway, B. R. (2021). Color space geometry uncovered with magnetoencephalography. Current Biology, 31(3), 515–526.

Rozman, A., Watts, D. J., Somers, L. P., Gunel, B., Racey, C., Barnes, K., & Bosten, J. M. (2025). Tuning of cortical color mechanisms revealed using steady-state visually evoked potentials. Imaging Neuroscience, 3, IMAG-a.

Shapley, R., Nunez, V., & Gordon, J. (2019). Cortical double-opponent cells and human color perception. Current Opinion in Behavioral Sciences, 30, 1–7.

Shapley, R., Nunez, V., & Gordon, J. (2025). Cortical processing of color: Chromatic visual evoked potentials. Vision research, 229, 108564.

Solomon SG, Lennie P. 2005. Chromatic gain controls in visual cortical neurons. J. Neurosci. 25:4779–92

Stockman, A. (2019). Cone fundamentals and CIE standards. Current Opinion in Behavioral Sciences, 30, 87–93.

Switkes, E., Crognale, M. A. (1999). Comparison of color and luminance contrast: apples versus oranges? Vision Research, 39(10), 1823–183. 10.1016/S0042-6989(98)00219-3

Tailby, C., Solomon, S. G., Dhruv, N. T., Majaj, N. J., & Lennie, P. (2005). Habituation reveals cardinal chromatic mechanisms in striate cortex of macaque. Journal of Vision, 5(8), 80–80.

Victor, J. D., Maiese, K., Shapley, R., Sidtis, J., & Gazzaniga, M. S. (1989). Acquired central dyschromatopsia: analysis of a case with preservation of color discrimination. Clinical Vision Sciences, 4(3), 183–196.

von der Twer, T., & MacLeod, D. I. (2001). Optimal nonlinear codes for theperception of natural colours. Network: Computation in Neural Systems, 12(3), 395.

Wang, J. and Wade, A.R. (2011). Differential attentional modulation of cortical responses to S-cone and luminance stimuli. Journal of Vision 11, 1. 10.1167/11.6.1

Webster, M. A. (2020). The Verriest Lecture: Adventures in blue and yellow. Journal of the Optical Society of America A, 37(4), V1–V14.

Webster, M. A., & Mollon, J. D. (1994). The influence of contrast adaptation on color appearance. Vision research, 34(15), 1993–2020.

Webster, M. A., & Wilson, J. A. (2000). Interactions between chromatic adaptation and contrast adaptation in color appearance. Vision research, 40(28), 3801–3816.

Winkler, A. D., Spillmann, L., Werner, J. S., & Webster, M. A. (2015). Asymmetries in blue–yellow color perception and in the color of ‘the dress’. Current Biology, 25(13), R547–R548.

Witzel, C., & Gegenfurtner, K. R. (2018). Color Perception: Objects, Constancy, and Categories. Annual Review of Vision Science, 4(1), 475–499. 10.1146/annurev-vision-091517-034231

Zaidi, Q., & Conway, B. (2019). Steps towards neural decoding of colors. Current Opinion in Behavioral Sciences, 30, 169–177.

